# Widespread dissemination of ESBL-producing *Salmonella enterica* serovar Infantis exhibiting intermediate fluoroquinolone resistance and harboring *bla_CTX-M-65_*-positive pESI-like megaplasmids in Chile

**DOI:** 10.1101/2023.09.25.559306

**Authors:** Alejandro Piña-Iturbe, Constanza Díaz-Gavidia, Francisca P. Álvarez, Rocio Barron-Montenegro, Diana M. Álvarez-Espejo, Patricia García, Doina Solís, Rodrigo Constenla-Albornoz, Magaly Toro, Jorge Olivares-Pacheco, Angélica Reyes-Jara, Jianghong Meng, Rebecca L. Bell, Andrea I. Moreno-Switt

## Abstract

**Background:** Multidrug-resistant (MDR) *Salmonella* Infantis has disseminated worldwide, mainly linked to the consumption of poultry products. Evidence shows dissemination of this pathogen in Chile; however, studies are primarily limited to phenotypic data or involve few isolates. As human cases of *Salmonella* Infantis infections have substantially increased in recent years, a better understanding of its molecular epidemiology and antimicrobial-resistance profiles are required to inform effective surveillance and control measures.

**Methods:** We sequenced 396 *Salmonella* Infantis genomes and analyzed them with all publicly available genomes of this pathogen from Chile (440 genomes in total), representing isolates from environmental, food, animal, and human sources obtained from 2009 to 2022. Based on bioinformatic and phenotypic methods, we assessed the population structure, dissemination among different niches, and AMR profiles of *Salmonella* Infantis in the country.

**Findings:** The genomic and phylogenetic analyses showed that *Salmonella* Infantis from Chile comprised several clusters of highly related isolates dominated by sequence type 32. The HC20_343 cluster grouped an important proportion of all isolates. The latter was the only cluster associated with pESI-like megaplasmids, and up to 12 acquired AMR genes/mutations predicted to result in an MDR phenotype. Accordingly, antimicrobial-susceptibility testing revealed a strong concordance between the AMR genetic determinants and their matching phenotypic expression, indicating that a significant proportion of HC20_343 isolates produce extended-spectrum β-lactamases and have intermediate fluoroquinolone resistance. HC20_343 *Salmonella* Infantis were spread among environmental, animal, food, and human niches, showing a close relationship between isolates from different years and sources, and a low intra-source genomic diversity.

**Interpretation:** Our findings show a widespread dissemination of MDR *Salmonella* Infantis from the HC20_343 cluster in Chile. The high proportion of isolates with resistance to first-line antibiotics and the evidence of active transmission between the environment, animals, food, and humans highlight the urgency of improved surveillance and control measures in the country. As HC20_343 isolates predominate in the Americas, our results suggest a high prevalence of ESBL-producing *Salmonella* Infantis with intermediate fluoroquinolone resistance in the continent.

**Funding:** Agencia de Investigación y Desarrollo de Chile (ANID) through FONDECYT de Postdoctorado Folio 3230796 and Folio 3210317, FONDECYT Regular Folio 1231082, and ANID – Millennium Science Initiative Program – ICN2021_044.

**Research in context:** *Evidence before the study:* In the last decade, emergent multidrug-resistant *Salmonella* Infantis has spread worldwide, primarily linked to poultry product consumption. However, in most countries from the Americas Region, such as Chile, the extent of the dissemination of emergent *Salmonella* Infantis and its molecular epidemiology remains unknown. In May and September 2023, an online search was conducted using the Google engine and the PMC database with the terms “*Salmonella*,” “Infantis,” and “Chile,” with no language restrictions. We assessed the results to select those presenting antimicrobial resistance, epidemiologic, or genomic data directly associated with isolates from Chile (13 studies). The selected studies showed that the prevalence of *Salmonella* Infantis in poultry-meat production systems, its resistance to different antibiotics, and the number of human cases of infection caused by this serovar have increased since 2014-2016. However, these reports were limited to phenotypic data or involved the genomic analysis of a few isolates (<50) obtained from the same source. No study has assessed the genomic epidemiology of the *Salmonella* Infantis population at the country level.

*Added value of this study:* Here, we present the first large-scale genomic epidemiology analysis of Salmonella Infantis in Chile, including isolates from environmental, food, animal, and human sources obtained from 2009 to 2022. We found that Salmonella Infantis in Chile is divided into several clusters of highly related isolates and that only a single cluster, the HC20_343, was associated with multiple antimicrobial-resistance determinants and pESI-like megaplasmids. We also report that isolates from this cluster are widespread among most sources, including irrigation water, poultry, food, and human cases. Detection of AMR determinants coupled with antimicrobial-susceptibility testing indicated that most HC20_343 isolates are ESBL-producers and have intermediate resistance to ciprofloxacin. Population structure analysis of this foodborne pathogen evidenced an active transmission of MDR Salmonella Infantis between different niches. This study reveals the widespread dissemination of MDR Salmonella Infantis in Chile.

*Implications of all the available evidence:* The evidence indicates that emerging *Salmonella* Infantis from the HC20_343 cluster is spreading among various niches, including irrigation water, poultry, and food, causing human infections in Chile. Its resistance to first-line antibiotics used for treating salmonellosis in individuals with a higher risk of severe or invasive infections is concerning. Currently, most surveillance and control efforts to reduce salmonellosis in Chile are focused on the poultry industry, and the study of outbreaks does not include whole-genome sequence analyses. Our findings highlight the urgent necessity to improve the surveillance and control measures to include agricultural waters to prevent contamination of produce and the further dissemination of resistance genes in the environment. As the HC20_343 cluster is highly prevalent in the Americas, further research involving large-scale genomic population analyses would shed light on the extent of the dissemination and transmission routes of emergent *Salmonella* Infantis in the continent and may contribute to informing surveillance and control policies.

## Introduction

Non-typhoidal *Salmonella* (NTS) is one of the leading causes of foodborne disease globally, mainly affecting children under five and the elderly. In 2019, NTS caused 215,000 deaths and the loss of approximately 15 million years of healthy life worldwide, according to the estimates of the Global Burden of Disease Study^1^ estimates. The burden posed by NTS is in part fueled by the emergence and spread of antimicrobial resistance, especially to critically-relevant antimicrobials such as third-generation cephalosporins and fluoroquinolones, which are the first-line treatment option for severe NTS infections^2,3^.

In the past decade, multidrug-resistant (MDR) and extended-spectrum β-lactamase (ESBL)-producing *Salmonella enterica* serovar Infantis have emerged in different continents as a zoonotic pathogen causing outbreaks of foodborne illness associated with poultry products consumption^4–6^. In some countries, this pathogen has displaced the historically most prevalent *Salmonella* serovars, such as Typhimurium and Enteritidis^4,7^. The success of emerging MDR *Salmonella* Infantis is linked to the acquisition of a ≈300 kbp pESI-like megaplasmid, which encodes virulence, fitness-enhancing, and antibiotic-resistance factors that favor its capacity of biofilm production, adhesion, invasion, and resistance to third-generation cephalosporins^4–6^. Moreover, isolates of this emerging pathogen are also associated with the chromosomal *gyrA* (D87Y) mutation, involved in fluoroquinolone resistance^5,8^. The enhanced virulence and antimicrobial resistance traits of emergent *Salmonella* Infantis make this pathogen a global threat to public health.

Chile is located between the Andes mountains and the Pacific Ocean, in the southernmost part of South America. Organized into 16 administrative Regions, Chile concentrates its population and most agricultural activities in the central area^9^, being one of the leading exporters of poultry meat in the region, ranking third after Brazil and the United States^10^. Recent research, primarily phenotypic or involving few isolates, has shown the spread of MDR *Salmonella* Infantis in the country, mainly linked to poultry. Analysis of the poultry and pig production systems indicated increased *Salmonella* Infantis prevalence and resistance to multiple antibiotics such as β-lactams, aminoglycosides, and tetracyclines^11,12^. This was concurrent with a high proportion of reported ESBL-producing isolates in poultry products sold at Santiago de Chile’s supermarkets^13–15^. In 2019, the Public Health Institute of Chile (ISP) reported a 431% increase in intestinal and invasive human infections caused by *Salmonella* Infantis in 2018, compared to 2014^16^, clearly documenting the farm-to-fork transmission of this pathogen. Furthermore, MDR strains of this pathogen also were isolated from irrigation water and a Magellanic Horned-Owl^17,18^, suggesting its dissemination in the environment outside poultry sources. These data document the emergence and spread of this foodborne pathogen in Chile in recent years; however, its magnitude and molecular epidemiology remain unknown.

In this study, we report the first large-scale genomic analysis of the *Salmonella* Infantis population in the country, assessing its population structure, dissemination among different sources, and antimicrobial resistance. This effort will produce valuable information for the entities responsible for the country’s surveillance and control of this pathogen. We sequenced 396 *Salmonella* Infantis genomes and analyzed them with all other available *Salmonella* Infantis genomes from Chile (440 genomes; April 20^th^, 2023), representing environmental, food, animal, and clinical strains. Our analysis revealed that emergent *Salmonella* Infantis, with resistance to first-line antibiotics, has been in Chile since before 2016, actively transmitted between environmental, animal, food, and human sources. These results make an urgent call to enhance surveillance and control measures to prevent the further spread of this pathogen and antibiotic resistance genes in food and the environment.

## Methods

### Sample collection and isolation of *Salmonella* Infantis

From 2014-2022, samples from environmental, poultry, food, animal feed, animal, and clinical sources were collected from three regions of central Chile: Región de Valparaíso, Región Metropolitana, and Región del Maule (**Fig. 1A** and **Supplementary Table S1**). Surface water samples (10 L) were collected at various points from five watersheds in central Chile (the Maipo, Mapocho, Claro, Lontué, and Mataquito rivers) using modified Moore swabs^19^. Poultry production samples (boot swabs, chicken crops, and cecal content) were collected from poultry farms and chicken-meat production systems located in Región Metropolitana. These samples were processed according to a modified FDA-BAM protocol to isolate *Salmonella* as previously described^9^. Raw meat-based dog diets and fecal samples from raw-fed dogs were processed according to the FDA-BAM protocol with modifications; raw food (25g) was enriched in 225 mL lactose broth (BD Difco), and fecal samples were enriched in 10 mL buffered peptone water.

**Figure 1.**
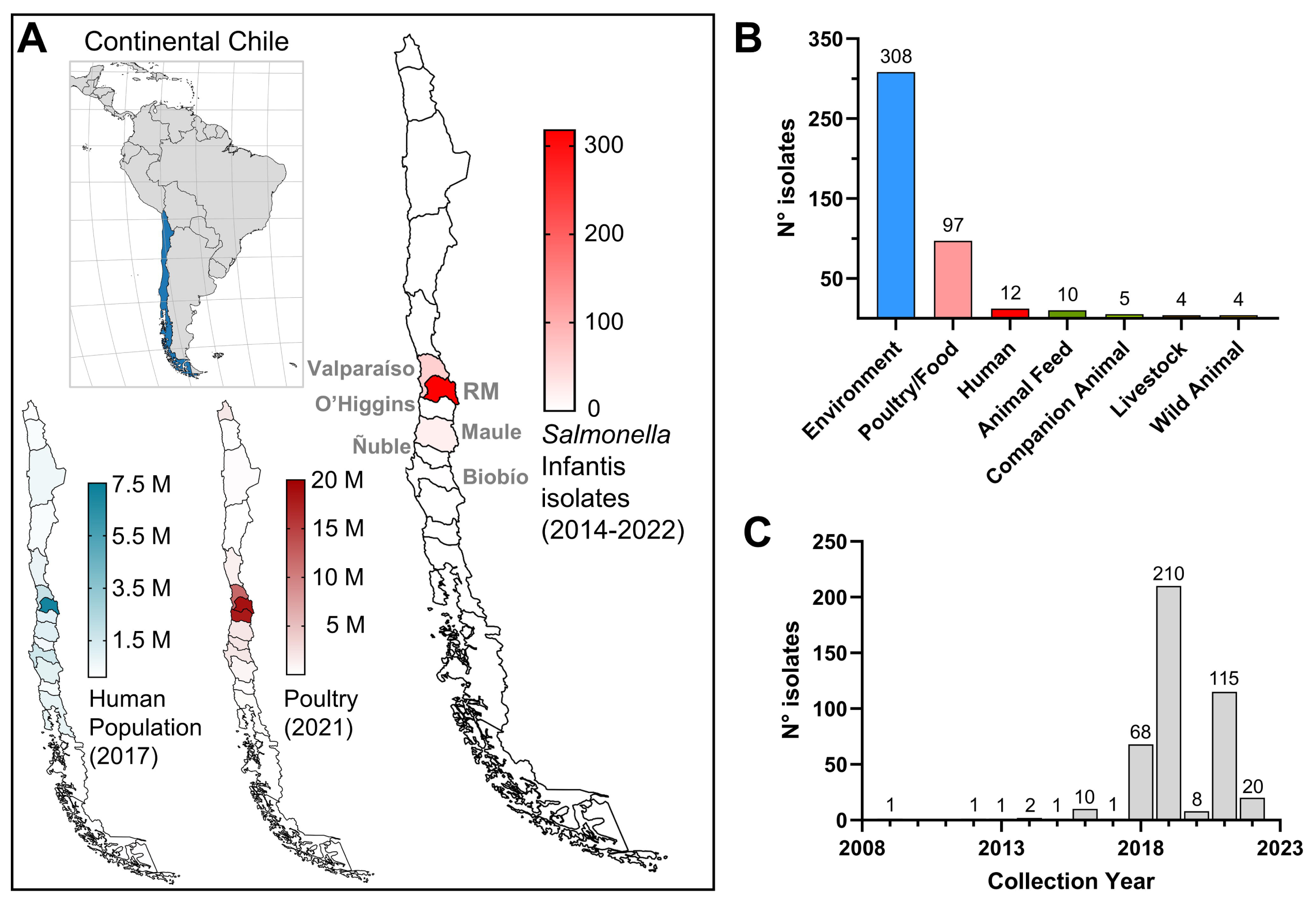
Region of origin, isolation source, and collection year of the *Salmonella* Infantis isolates analyzed in this study. **A**) Location of continental Chile in South America and distribution of its human population, poultry production, and collected *Salmonella* Infantis isolates among the 16 Regions (first-level administrative divisions). Note the concentration of these three variables in central Chile, especially in the Metropolitan Region. Population and poultry data were obtained from the corresponding last population and agricultural censuses carried out in 2017 and 2021, respectively (available at http://resultados.censo2017.cl/ and https://www.ine.gob.cl/censoagropecuario/resultados-finales/graficas-regionales. The administrative Regions from which the *Salmonella* Infantis isolates were collected are indicated. RM: Región Metropolitana. **B**) *Salmonella* Infantis isolates distribution per isolation source and per **C**) collection year.

Isolation of *Salmonella* was confirmed by PCR amplification of the *invA* gene with primers *invA*F (5′-GAATCCTCAGTTTTTCAACGTTTC-3′) and *invA*R (5′- TAGCCGTAACAACCAATACAAATG-3′)^20^.

Raw and ready-to-eat poultry products were collected from supermarkets, restaurants, and meat-producer facilities by the Subsecretaría de Salud Pública from Valparaíso and transported at 0-4°C to the Laboratorio de Salud Pública, Ambiental y Laboral (SEREMI Salud – Valparaíso). *Salmonella* isolation from these samples was performed according to ISO 6579-2017, and serotyping was performed at Instituto de Salud Pública de Chile (ISP; Public Health Institute of Chile).

Human clinical samples (e.g., stool, blood, urine) representing human salmonellosis cases from different Región Metropolitana areas were received at the Laboratory of Microbiology of the UC-Christus Health network and processed for *Salmonella* isolation. Samples were inoculated in Hektoen Enteric agar and incubated at 35±2°C for 24-48 hours in aerobiosis; then, *Salmonella* confirmation was performed using MALDI-TOF mass spectrometry, and serotyping was performed at ISP, Chile.

*Salmonella* Infantis available from the above-described sources were all selected. All isolates were stored in 20% glycerol stocks and maintained at -80°C.

### Genome sequencing and construction of the genome dataset

Isolates from surface water were sequenced at the Food and Drug Administration, Center for Food Safety and Applied Nutrition. Isolates from other sources were sequenced at the GenomeTrakr New York State Department of Health laboratory. Whole genome sequencing was performed on 385 isolates, and the reads were deposited in the Sequence Read Archive, NCBI. Additionally, the *Salmonella* Infantis strains from human cases were sequenced at SeqCenter, Pittsburgh, PA, and the reads were directly uploaded to Enterobase^21^. All sequencing was conducted using Illumina platforms.

On April 20^th^, 2023, the Enterobase database for *Salmonella* was queried for genomes from Chile and the serovar Infantis as predicted by SISTR1^22^ or SeqSero2^23^. A total of 440 *Salmonella* Infantis records were retrieved, including the 396 sequenced by us, along with their associated metadata (**Table 1**; **Supplementary Table S1**). All genome assemblies were downloaded from Enterobase.

**Table 1.** Features of the publicly available *Salmonella* Infantis genomes from Chile (April 20^th^, 2023)

| Feature | # genomes (Total = 440) |
| --- | --- |
| 7-gene MLST |  |
| ST32 (*thrA19*) | 435 (98.9%) |
| ST9835 (*thrA1541*) | 3 (0.7%) |
| ST9853 (*thrA1552*) | 2 (0.5%) |
| cgMLST + HierCC |  |
| HC20_343 | 322 (73.2%) |
| HC20_775 | 83 (18.9%) |
| HC20_2398 | 8 (1.8%) |
| other HC20 clusters | 27 (6.1%) |
| Isolation source |  |
| Environment<sup>a</sup> | 308 (70.0%) |
| Food/Poultry | 97 (22.0%) |
| Human | 12 (2.7%) |
| Other | 23 (5.2%) |
| Isolation year |  |
| 2022 | 20 (4.5%) |
| 2021 | 115 (26.1%) |
| 2020 | 8 (1.8%) |
| 2019 | 210 (47.7%) |
| 2018 | 68 (15.5%) |
| 2009-2017 | 19 (4.3%) |

<sup>a</sup>Environmental isolates were obtained from surface water (n=265) and soil/dust (n=43).

### Population structure and phylogenetic analyses

The 7-gene MLST, core genome MLST (cgMLST), hierarchical clustering based on cgMLST profiles, core SNP-based phylogeny, and minimum spanning trees were all carried out directly in Enterobase^21^. Annotated genomes were used for allele calling to classify the *Salmonella* Infantis isolates in sequence types (STs) by MLST (based on genes *aroC*, *dnaN*, *hemD*, *hisD*, *purE*, *sucA* and *thrA*) or core genome STs (cgSTs) by cgMLST (based on a 3002 allele scheme, cgMLST V2; https://pubmlst.org/bigsdb?db=pubmlst_salmonella_seqdef&page=schemeInfo&scheme_id=4). HierCC V1^24^ was used to cluster isolates based on the cgMLST profiles. Clusters grouping isolates with links no more than 20 alleles apart (HC20) were used to describe the *Salmonella* Infantis population. Based on cgMLST profiles, a minimum spanning tree was constructed with MSTree V2 and visualized with GrapeTree v1.5.1^25^. A maximum likelihood phylogenetic tree based on core-SNPs was built with the *Salmonella* Infantis N55391 genome (Enterobase Barcode SAL_EA1888AA; GenBank accession NZ_CP016410.1) as a reference. The phylogenetic tree was constructed with RAxML V8 based on 2008 variant sites called in ≥95% of the genomes. A detailed description of how the Enterobase processes and analyses work is available in reference ^21^ and its supplementary files. The iTOL v6 online tool was used to display and annotate the phylogenetic tree (https://itol.embl.de/).

### Identification of antibiotic-resistance genes/mutations and presence of pESI- like megaplasmids

All 440 genome assemblies were downloaded from Enterobase and stored locally. Antibiotic resistance genes and point mutations involved in antimicrobial resistance in *Salmonella* were identified using AMRFinderPlus v3.11.4^26^. Since the *mdsA* and *mdsB* genes were found in all isolates and their presence did not result or explain any phenotypic resistance, these genes were not included in the analyses. The presence of pESI-like megaplasmids was assessed with ABRicate v1.0.1 (https://github.com/tseemann/abricate) and a custom database that included all 315 genes from the pESI-like megaplasmid pN55391 (GenBank accession NZ_CP016411.1).

### Intra-source genomic diversity analysis

The cgMLST allelic profiles for each *Salmonella* Infantis isolate were downloaded from Enterobase. Pairwise allelic distances (PAD) were calculated for any pair of isolates within each source to assess the genomic diversity within each source.

Only isolates representing unique cgSTs within a given source were included in the analysis. When more than one isolate per cgST was detected, only one was randomly selected and included in the analysis.

### Antibiotic-susceptibility testing

A sub-sample of 23 *Salmonella* Infantis isolates representing the diversity of the HC20_343 cluster was selected for antimicrobial susceptibility testing. These isolates were chosen because they represented different isolation sources and collection years and had the highest intra-source genomic diversity based on their PADs. Susceptibility testing was carried out in cation-adjusted Müeller-Hinton agar by the agar-dilution method following the recommendations of the Clinical and Laboratory Standards Institute^27^. The minimum inhibitory concentration (MIC) was interpreted according to the CLSI breakpoints available in the M100Ed33 document^28^. The following antibiotics, or antibiotic-inhibitor, were tested: amikacin (AMK), ampicillin (AMP), ampicillin-sulbactam (SAM), cefazoline (CFZ), cefepime (FEP), cefotaxime (CTX), cefotaxime/clavulanate (CTX/CLA), ceftazidime (CAZ), ceftazidime/clavulanate (CAZ/CLA), ciprofloxacin (CIP), fosfomycin (FOS), gentamycin (GEN), imipenem (IPM), meropenem (MEM), piperacillin-tazobactam (TZP), and trimethoprim/sulfamethoxazole (SXT). ESBL production was detected when the MIC of CTX and CAZ showed ≥3 2-fold reduction in the presence of clavulanate, an ESBL-inhibitor. A multidrug-resistant phenotype was assigned to isolates that displayed resistance to one or more antibiotics from at least three different classes.

## Results

### Region of origin, isolation source, and collection year of the isolates

To study the structure of the *Salmonella* Infantis population from Chile, we sequenced 396 isolates of this pathogen. We analyzed their genomes with all public *Salmonella* Infantis genomes from Chile available in Enterobase on April 20^th^, 2023. Our dataset was mainly composed of genomes from isolates coming from central Chile, specifically from Región de Valparaíso (n=60), Región Metropolitana (n=378), Región del Libertador General Bernardo O’Higgins (n=1), Región del Maule (n=21), Región Ñuble (n=1), and Región del Biobío (n=2). Most isolates (85%) came from Región Metropolitana and Región de Valparaíso (**Fig. 1A**). Genomes from environmental (river/creek/lagoon/irrigation water; boot swabs of soil/dust) and poultry/food (chicken carcass/crop/cecal content/feces; chicken meat) origin made 91% of total genomes, followed by human clinical cases which accounted for 2.7% (12 genomes) (**Fig. 1B**). While the collection year ranged from 2009 to 2022, most isolates (421/95.7%) were distributed from 2018 to 2022 (**Fig. 1C**). Although this dataset is limited to 6 out of 16 Regions in Chile and is primarily concentrated in two regions, it is relevant to note that these regions harbor most of the human population and poultry production in Chile (**Fig. 1A**). Therefore, the genome dataset analyzed in this study offers a substantial representation of the bacterial population potentially encountered by most individuals and poultry raising/production systems in the country.

### The *Salmonella* Infantis population in Chile belongs to ST32 and comprises different cgMLST clusters of highly related isolates

Among all isolates, 435 (98.9%) were ST32 (7-gene MLST) while the other five (three ST9835 and two ST9853) were single locus variants of ST32 that differed in the *thrA* allele (**Table 1**). The core SNP-based maximum likelihood phylogeny showed the presence of several clades comprising highly related genomes (**Fig. 2**). The hierarchical clustering of genomes linked by no more than 20 cgMLST-allele differences (HC20) revealed that the clades shown by the phylogeny corresponded with different HC20 groups, dominated by clusters HC20_343 and HC20_775 (92% of genomes) (**Table 1**, **Fig. 2**). Isolates belonging to cluster HC20_343 came from the highest diversity of isolation sources and some clusters (e.g., 2398, 422, 215390, 327110) seemed to group isolates only from environmental sources (**Fig. 2**). However, this is most likely an effect of a sample-size bias, with the most populated cluster being more likely to harbor genomes from different sources. Notably, 10 out of 12 *Salmonella* Infantis isolates from human infections belonged to the HC20_343 cluster. These isolates were closely related to isolates from different sources, forming subclades in the phylogeny that included poultry (chicken), food (chicken meat), and environmental (surface water and soil/dust from boot swabs) isolates (**Fig. 2**). The interspersed distribution of isolates from diverse sources across the phylogenetic tree that share temporal proximity (sampled during 2020-2022) suggests an extant transmission network involving environmental, animal, food, and human niches.

**Figure 2.**
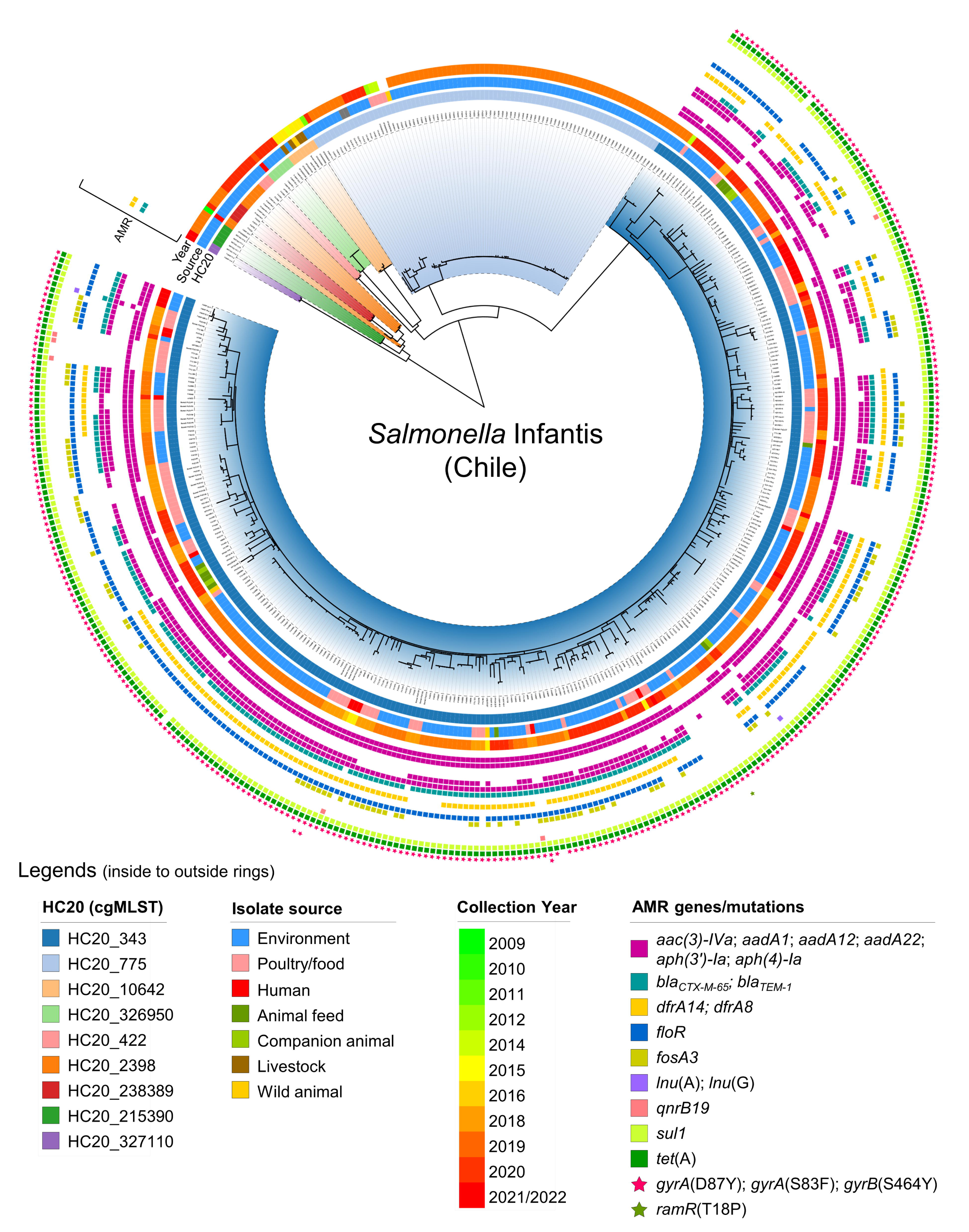
Phylogenetic analysis of Chilean *Salmonella* Infantis genomes. A core SNP-based maximum likelihood phylogeny (2008 variant sites; 95% presence) was constructed with 440 Chilean *Salmonella* Infantis genomes using *Salmonella* Infantis N55391 (Enterobase barcode SAL_EA1888AA), a USA strain isolated from poultry in 2014, as the reference. Additionally, metadata regarding the HC20 clusters (clade colors and first ring), isolation source (second ring), isolation year (third ring), and presence of antibiotic-resistance genes/mutations as determined by AMRFinderPlus (colored squares/stars) were incorporated into the phylogenetic tree.

### *Salmonella* Infantis isolates from the HC20_343 cluster include MDR ESBL-producing strains and carry pESI-like megaplasmids

The presence of antimicrobial-resistance (AMR) genes/mutations among the 440 genomes from Chile was assessed (**Fig. 2**; **Supplementary Table S2-S3**).

Interestingly, the HC20_343 cluster (322 genomes) was the only one associated with up to 12 acquired antimicrobial resistance genes or mutations predicted to result in resistance to aminoglycosides, cephems, folate pathway antagonists, chloramphenicol, fosfomycin, lincosamides, fluoroquinolones and tetracycline (**Fig. 3A**). The most frequent resistance genes among HC20_343 isolates were *tet*(A) (99.4%), *sul1* (99.4%), and *aadA1* (98.8%), encoding predicted resistance to tetracycline, sulfonamide, and aminoglycosides. Genes *aph(3’)-Ia*, *aph(4)-Ia*, *aac(3)-IVa* (aminoglycoside resistance), *bla_CTX_*_-M-65_ (third-generation cephalosporin resistance), *dfrA14* (trimethoprim resistance), and *floR* (phenicol resistance) were found in 60.6%-81.7% of the isolates. Other identified resistance genes [*aadA12*, *aadA22*, *bla_TEM-1_*, *lnu*(A), *lnu*(G), and *qnrB19*] were present in 6 or less isolates only, except by *fosA3* (fosfomycin resistance) that was carried by 28.9% of isolates. Noteworthy, all but one HC20_343 isolate (99.7%) carried the *gyrA* (D87Y) mutation involved in fluoroquinolone resistance.

**Figure 3.**
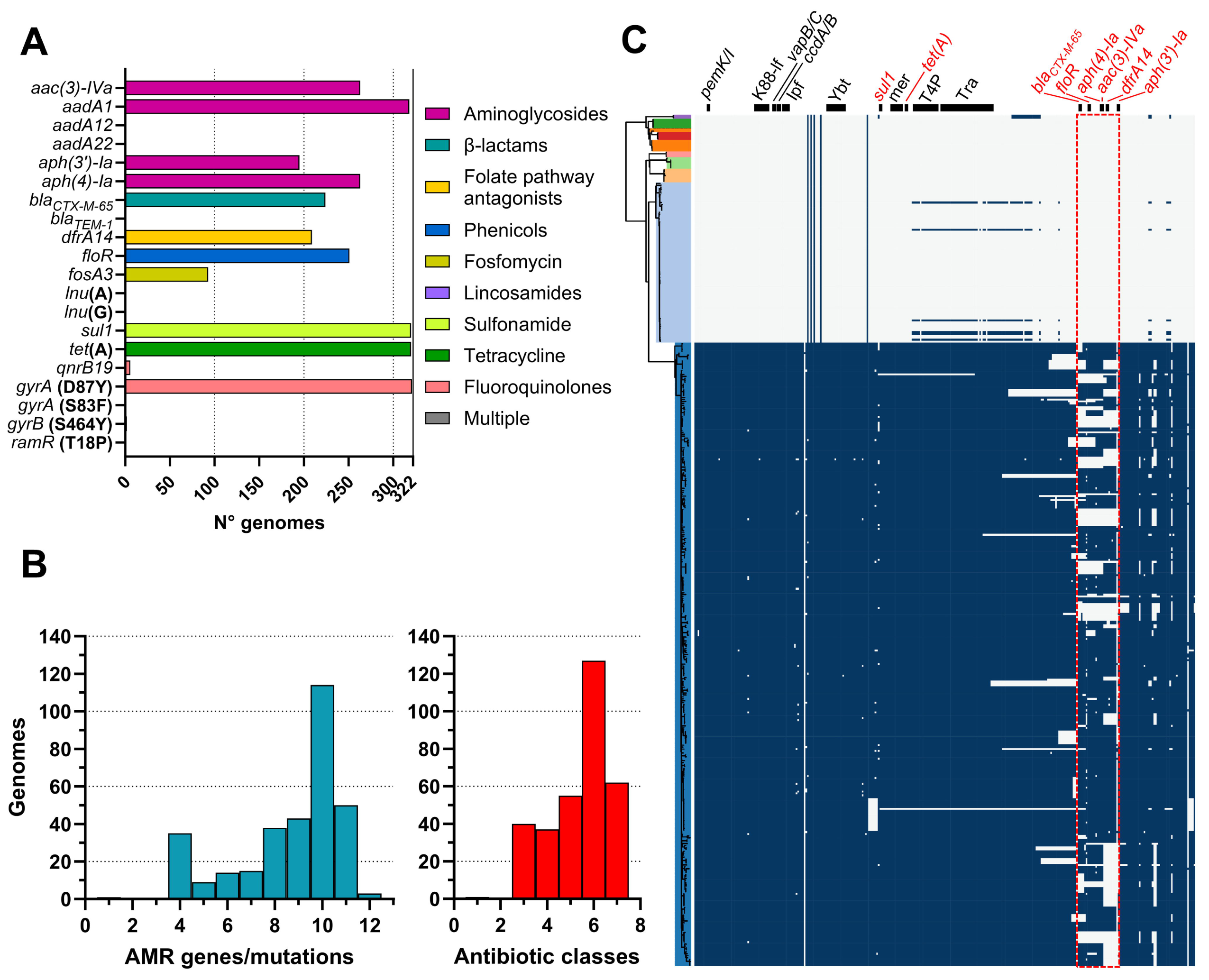
Antibiotic-resistance determinants and pESI-like megaplasmid presence among *Salmonella* Infantis from the HC20_343 cluster. **A**) Frequency of individual antibiotic-resistant genes/mutations in the HC20_343 colored by antibiotic class. **B**) Frequency of overall antibiotic-resistance genes/mutations per genome (blue bars) and antibiotic classes targeted per genome (red bars). **C**) Presence of pESI-like megaplasmids among the Chilean *Salmonella* Infantis genomes. A vertical representation of the same phylogenetic tree of Fig. 2 is colored according to the HC20 clusters. The presence of each of the 315 genes from the pESI-like megaplasmid pN55391 was assessed by ABRicate using a custom database (blue squares). Black horizontal bars above the presence/absence matrix indicate the backbone regions of the megaplasmid (genes in black font) or the antibiotic-resistance regions (genes in red font). A red dashed line rectangle delimits the most variable region among the Chilean pESI- like megaplasmids relative to pN55391. K88-lf (K88-like fimbria), Ipf (Infantis plasmid-encoded fimbria, Ybt (Yersiniabactin synthesis cluster), mer (mercury- resistance cluster), T4P (type-IV pili encoding cluster), Tra (transfer region).

Within the HC20_343 cluster, 210 genomes (65.2%) harbored nine or more AMR genes/mutations, and 321 genomes (99.7%) carried genes/mutations predicted to encode resistance to at least one antibiotic from three or more antibiotic classes, potentially resulting in a multidrug-resistant phenotype (**Fig. 3B**). We found a good agreement between the content of genetic AMR determinants and the phenotypic resistance in all the isolates for which the MIC was assessed (**Supplementary Table S4**). Accordingly, all *bla_CTX_*_-M-65_-positive isolates were resistant to the cephalosporins CFZ and CTX and displayed an ESBL-phenotype. Isolates harboring *aadA1*, *aac(3)-Iva,* and *aph(4)-Ia* were resistant to GEN. Many isolates (9/23) had intermediate susceptibility to the fourth-generation cephalosporin FEP. However, this phenotype did not correlate with any of the identified AMR genes. The presence of *aadA1* alone was not sufficient to confer GEN resistance, and *aph(3’)-Ia* carriage did not show agreement with the resistance profile. All tested isolates were susceptible to AMK. SXT-resistance was found in isolates carrying *sul1* plus *dfrA14*, and lack of *dfrA14* resulted in SXT susceptibility. All *gyrA*(D87Y)- positive isolates displayed intermediate resistance to CIP. Notably, the presence of *qnrB19* plus *gyrA*(D87Y) resulted in CIP-resistance. Conversely, all isolates lacking at least one of the genes/mutations mentioned above were susceptible to the corresponding antibiotics.

Since many of the identified antibiotic-resistance genes have been reported to be carried by the pESI-like megaplasmid associated with emerging MDR *Salmonella* Infantis, we screened the entire genome dataset for the presence of the pN55391 pESI-like megaplasmid genes (**Fig. 3C**; **Supplementary Table S5**). We found that only the HC20_343 cluster harbored most of the pESI-like megaplasmid genes (from 164 to 310 out of 315 genes), including those encoding the three toxin-antitoxin systems (*pemK/I*, *vapB/C* and *ccdA/B*), the K88-like and Ipf fimbria, the yersiniabactin synthesis cluster, the mercury resistance cluster, and the conjugative transfer region (*tra* and type-IV pili-encoding genes), which are part of the pESI-like megaplasmids backbone^29^. Most differences between the megaplasmids harbored by the Chilean strains and the pN55391 megaplasmid were located in the antibiotic resistance region, previously reported as a variable region ^5^. Nevertheless, the resistance genes contained in this region were present in most of the megaplasmid-harboring genomes (*bla_CTX-M-65_*: 69.6%; *floR*: 78.0%; *aph(4)-Ia*: 81.7%; *aac(3)-IVa*: 81.7%; *dfrA14*: 64.9%; and *aph(3’)-Ia*: 60.6%; **Supplementary Table S5**). Our analyses revealed that the HC20_343 *Salmonella* Infantis isolates acquired multiple antimicrobial-resistance genes/mutations, partly associated with the presence of pESI-like megaplasmids.

### HC20_343 *Salmonella* Infantis isolates are disseminated among different niches and show a low intra-source genomic diversity

A minimum spanning tree (MST) was constructed to visualize the genomic structure of the *Salmonella* Infantis population based on the cgMLST profiles (**Fig. 4A**). In agreement with the core SNP-phylogeny, the MST shows the bacterial population grouped in nine HC20 clusters of highly related isolates linked by no more than 20 allele-differences. The highest diversity of sources was found for cluster HC20_343, while the other HC20 clusters came mainly from environmental samples. Only two out of 12 isolates from human clinical samples were found outside HC20_343 in clusters HC20_215390 and HC20_2398. All isolates from food, animal feed, and companion animals (dogs) belonged to cluster HC20_343. These findings highlight the foodborne, zoonotic, and human-pathogenic potential of HC20_343 *Salmonella* Infantis.

**Figure 4.**
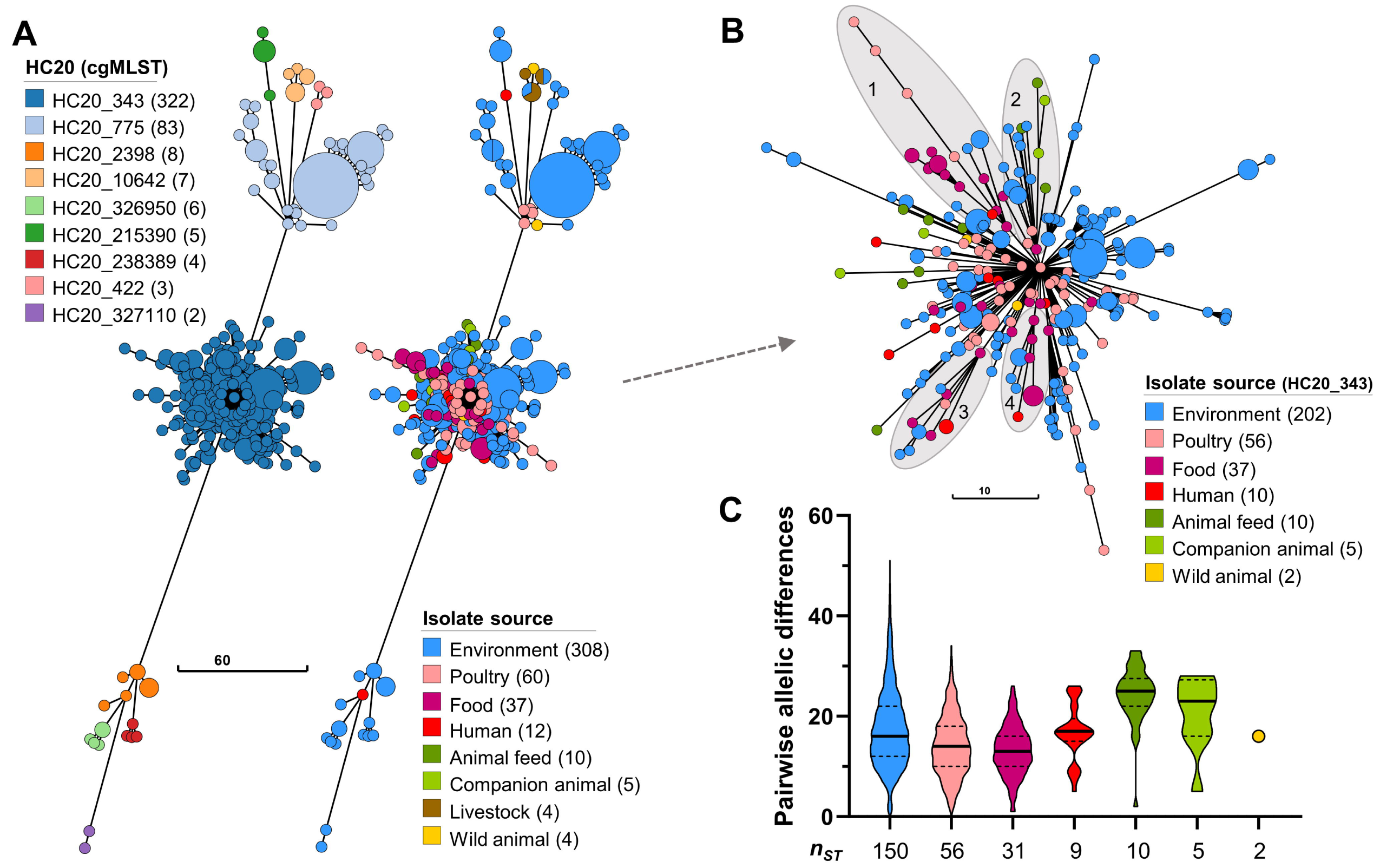
Population structure of Chilean *Salmonella* Infantis and intra- source genomic diversity within cluster HC20_343. **A**) Minimum spanning tree depicting the population structure of *Salmonella* Infantis in Chile based on the cgMLST profiles of 3002 alleles. Nodes are colored according to the HC20 clusters determined by HierCC or the isolation source, and their size is proportional to the number of isolates included in each node. The legends indicate the different HC20 clusters present, the isolation sources, and the number of isolates. **B**) Zoom-in view of the HC20_343 cluster structure with nodes colored according to the isolation source. The legend also indicates the number of isolates per source within HC20_343. The shaded area and numbers indicate subclusters evidencing transmission of *Salmonella* Infantis between different sources. **C**) Pairwise allelic differences between unique cgSTs from cluster HC20_343 per isolation source. Each violin plot, truncated at the highest and smallest values, represents the frequency distribution of PADs. The black unbroken and dashed lines represent the median PAD, and the 25^th^ and 75^th^ percentiles, respectively. The ***n_ST_*** value indicates the number of isolates with unique cgSTs within each isolation source. Only isolates representing unique cgSTs were included in the analysis.

We carried out a more detailed analysis of cluster HC20_343 isolates. The MST evidenced putative events of transmission between different sources, as exemplified by subclusters 1 to 4 in **Fig. 4** **B**. Subclusters 1, 3, and 4 included isolates from environmental, food, and human sources, while subcluster 2 included isolates from environmental, animal feed, and companion animals. Moreover, subclusters 1, 2, and 3 also included *Salmonella* Infantis from poultry. Importantly, isolates from all sources within these clusters were linked to isolates obtained from food (poultry products), and, ultimately, all sources within cluster HC20_343 had links with poultry. We assessed the intra-source genomic diversity of the HC20_343 isolates regarding the pairwise allelic differences (PAD) between any pair of isolates representing unique cgSTs (**Fig. 4C**; **Supplementary Table S6**).

Overall, a low genomic diversity was found within the different isolation sources. The environmental isolates displayed the highest diversity, with PAD values ranging from 0 to 51. All other sources harbored isolates with PADs ≤34. The median PAD per source ranged from 13 in food isolates to 25 in animal feed.

Environmental, poultry, food, and human isolates had the lowest median PADs (from 13 to 17), while the isolates from animal feed and companion animals had the highest (25 and 23, respectively). Overall, the close relatedness between HC20_343 isolates from different sources and the low intra-source genomic diversity further supports a scenario in which MDR *Salmonella* Infantis is actively disseminating among different niches, including humans, in Chile.

## Discussion

The expansion of MDR *Salmonella* Infantis has been reported worldwide, with the highest proportion of isolates coming from the Americas region, followed by Europe^30^. The dissemination of this foodborne pathogen is mainly linked to poultry and poultry products, and different countries have reported a rise in human infections^7,12,30–32^. Nevertheless, little is known about the extent of the *Salmonella* Infantis dissemination in different niches and its molecular epidemiology.

Here, we reported the first large-scale genomic analysis of this foodborne pathogen population in Chile, finding evidence of the transmission of *Salmonella* Infantis carrying pESI-like megaplasmids and multiple AMR determinants (cluster HC20_343) between environmental, poultry, food, animals, and human niches.

Highly related isolates from different years were found in diverse sources, indicating a constant inter-source transmission. The country is working to enhance the surveillance and control of *Salmonella* (including antimicrobial resistance) (SAG, Exempt Resolution 3687; https://bcn.cl/2pap1)^13^. However, these efforts mainly focus on poultry and its derived products as they are known sources of *Salmonella*. Importantly, we report that irrigation waters are a source of *Salmonella* Infantis with potential MDR phenotypes. The presence of this pathogen in surface watersheds that supply the country’s main agricultural region^9^ underscores the urgent necessity to improve the current monitoring of irrigation waters and establish effective control measures to prevent the contamination of produce and the dissemination of antibiotic-resistance genes in the environment.

Human infection with non-typhoidal *Salmonella* usually results in self-limited acute gastroenteritis. However, children under five, adults over 65, and immunocompromised people are at higher risk of developing a severe life- threatening infection^3^. Antibiotics, such as third-generation cephalosporins and fluoroquinolones, are recommended to prevent or treat severe diseases^3^. In 2017, the World Health Organization presented a list of 12 antibiotic-resistant bacterial pathogens, categorized into critical, high, and medium priority tiers, urgently requiring research and development of novel antibiotics since available treatments are becoming limited^33^. This list placed third-generation cephalosporin-resistant *Enterobacteriaceae* and fluoroquinolone-resistant *Salmonella* spp. as critical and high-priority pathogens. A significant proportion of *Salmonella* Infantis from the HC20_343 cluster (69.6%) harbored the ESBL-encoding gene *bla_CTX-M-65_* in a pESI-like megaplasmid. Moreover, almost all HC20_343 isolates (321/322) harbored the chromosomal *gyrA* (D87Y) mutation involved in fluoroquinolone resistance.

Although the resistance profiles of emergent *Salmonella* Infantis reported in different countries are variable^7,31,34,35^, partially as a result of the diversity within the megaplasmid AMR region (see Fig. 3C and ref. ^5^), our findings in Chile are in agreement with a recent global survey of reported AMR determinants in *Salmonella* Infantis^30^. Importantly, we found that the aminoglycoside AMK and the carbapenems IPM and MEM were consistently active against all tested isolates from Chile. Our findings imply that the available options for preventing or treating severe infections in susceptible individuals are limited.

An unexpected finding was the association of the HC20_343 isolates with the presence of the pESI-like megaplasmids. Alba *et al*.^6^ found pESI-like megaplasmids carrying the *bla_CTX-M-1_* gene in European *Salmonella* Infantis isolates from a higher diversity of HC20 clusters, and mainly from the HC20_7898 cluster. They also reported that megaplasmid-positive isolates from North America, or associated with traveling to South America, harbored the *bla_CTX-M-65_*gene on the megaplasmid and belonged to the HC20_343 cluster. Similar to our study, they found that strains from the HC20_775 cluster lacked the megaplasmid. In addition to the known association of *bla_CTX-M-65_* with American and *bla_CTX-M-1_* with European megaplasmid-carrying *Salmonella* Infantis^6,7^, our findings suggest that the American megaplasmid-positive *Salmonella* Infantis strains might be associated with the HC20_343 cluster. Testing this hypothesis might help to better understand the global dissemination of emerging *Salmonella* Infantis. Importantly, we found that, out of 14706 *Salmonella* Infantis genomes from the Americas, 60% belonged to the HC20_343 cluster (**Supplementary File**, **FigS1**), suggesting a high prevalence of megaplasmid-positive ESBL-producing isolates with intermediate fluoroquinolone resistance in the continent.

The approximate time for the arrival of the emerging *Salmonella* Infantis into Chilean territory remains unknown. The oldest pESI-like positive genomes in our dataset date from 2016 (4 genomes) and 2017 (1 genome) (**Suplemental Table S1**). These genomes represent four clinical isolates and one isolate obtained from a Dominican gull (*Larus dominicanus*), indicating that *Salmonella* Infantis carrying *bla_CTX-M-65_*-positive pESI-like megaplasmids were circulating in Chile before 2016. A minimum spanning tree constructed with the cgMLST profiles from all *Salmonella* Infantis isolates from the Americas (available in Enterobase on July 7^th^, 2023) revealed that the HC20_343 isolates from the United States and Chile cluster together (**Supplementary File, FigS1,** also seen at **<u>NCBI Pathogen Detection</u>**). This finding suggests two possible scenarios in which emerging *Salmonella* Infantis from Chile came from the United States, or isolates from both countries share a common origin^36^.

We analyzed a dataset of 440 genomes representing the population structure of *Salmonella* Infantis in Chile. This population comprises strains lacking genetic determinants of antibiotic resistance and antibiotic-resistant strains that harbor pESI-like megaplasmids, both from the globally spread ST32. The megaplasmid-carrying strains belonged to the HC20_343 cluster, circulating among environmental, food, diverse animals, and human niches. Our results indicate that a significant proportion of the HC20_343 isolates encode ESBLs and display an intermediate resistance to fluoroquinolones, which limits the available treatments for individuals at a higher risk. Our findings and the reported increase in human cases highlight the urgent need to study the dissemination dynamics of this pathogen to devise effective surveillance and control measures.

## Contributors

Conceptualization - API, AIMS

Data curation - API, FPA, RBM, DMAE, PG, DS, JOP, ARJ, AIMS

Formal analysis - API, CDG, FPA, RBM, DMAE Funding acquisition - API, DMAE, ARJ, AIMS

Investigation - API, CDG, FPA, RBM, DMAE, PG, DS, RCA, MT, JOP, ARJ, JM, RLB, AIMS

Methodology - API, CDG, FPA, RBM, DMAE, PG, DS, RCA, MT, JOP, ARJ, JM, RLB, AIMS

Software - API, CDG Supervision - AIMS Visualization - API

Writing original draft - API, AIMS

Writing review and editing - API, CDG, FPA, RBM, DMAE, PG, DS, RCA, MT, JOP, ARJ, JM, RLB, AIMS

All authors read and approved the submitted version of the manuscript.

## Data sharing statement

All genome metadata, identified AMR genes/mutations, antimicrobial susceptibility testing results, and identified megaplasmid genes are available in the Supplementary Material. Genome assemblies are publicly available in Enterobase (https://enterobase.warwick.ac.uk/species/index/senterica) and GenBank (https://www.ncbi.nlm.nih.gov/genbank/) using the corresponding accession numbers found in the Supplementary Table S1.

## Acknowledgements

This work was supported by Agencia de Investigación y Desarrollo de Chile (ANID) through FONDECYT de Postdoctorado Folio 3230796 (to A.P-I.) and Folio 3210317 (to D.A-E), FONDECYT Regular Folio 1231082 (to A.I.M-S), and ANID –Millennium Science Initiative Program – ICN2021_044 (to A.R-J).The funders of the study had no role in study design, data collection, data analysis, data interpretation, or the writing of the report.

